# Cellular Synchronisation through Unidirectional and Phase-Gated Signalling

**DOI:** 10.1101/2020.11.26.399683

**Authors:** Gregory Roth, Georgios Misailidis, Charisios D. Tsiairis

**Author notes:** These authors contributed equally to this work (Gregory Roth, Georgios Misailidis).

## Abstract

Multiple natural and artificial oscillator systems achieve synchronisation when oscillators are coupled. The coupling mechanism, essentially the communication between oscillators, is often assumed to be continuous and bidirectional. However, the cells of the presomitic mesoderm synchronise their gene expression oscillations through Notch signalling, which is intermittent and directed from a ligand-presenting to a receptor-presenting cell. Motivated by this mode of communication we present a phase-gated and unidirectional coupling mechanism. We identify conditions under which it can successfully bring two or more oscillators to cycle in-phase. In the presomitic mesoderm we observed the oscillatory dynamics of two synchronizing cell populations and record one population halting its pace while the other keeps undisturbed, as would be predicted from our model. For the same system another important prediction, convergence to a specific range of phases upon synchronisation is also confirmed. Thus, the proposed mechanism accurately describes the coordinated oscillations of the presomitic mesoderm cells and provides an alternative framework for deciphering synchronisation.

---

Oscillator systems ranging from the Belousov-Zabotinsky chemical reactions(Ross et al., 1988) to humans clapping hands(Néda et al., 2000), reach synchrony when oscillators are coupled. Synchronisation is achieved when a common pace is established and the phase difference between oscillators is fixed. Under the most rigorous definition, the phase difference between oscillators is zero and they cycle in unison(Pikovsky et al., 2003). For two oscillators that are out of phase this can be achieved if the oscillator that is ahead in phase slightly decelerates while the lagging one slightly accelerates to catch up with the other one. The oscillators monitor constantly the phase difference between them and minimize it. A generic mathematical framework that captures these phenomena has been proposed by Kuramoto(Kuramoto, 2012; Strogatz, 2000) and is still widely applied to understand among others, the coordinated behaviour of cellular(Yoshioka-Kobayashi et al., 2020) and human networks(Saw et al., 2019; Shahal et al., 2020). Despite the mathematical elegance and broad applicability of this model, it makes specific assumptions about the symmetric behaviour of the oscillators, their continuous and bidirectional communication. The presomitic mesoderm (PSM) presents a synchronising system where Kuramoto coupling has been employed to explain their coordinated behaviour(Murray et al., 2011; Uriu et al., 2017; Venzin and Oates, 2020; Yoshioka-Kobayashi et al., 2020) although such underlying assumptions do not necessarily hold.

During vertebrate development the PSM cells build the somites and, to achieve that, they undergo synchronised genetic oscillations. More than one hundred genes are expressed in an oscillatory fashion with a period matching the one of somite formation(Dequeant et al., 2006). Synchronisation of the PSM cells’ activity depends on communication through Notch pathway(Jiang et al., 2000; Okubo et al., 2012; Riedel-Kruse et al., 2007). Due to cis-inhibition in Notch signalling, at any given time a cell can either have a surplus of the ligand Delta and send a signal, or of the receptor Notch and receive a signal(Sprinzak et al., 2010). Hence, this communication mechanism is unidirectional and has been described as analogous to “walkie/talkie”(Sprinzak et al., 2011). In addition, the ligand Delta is periodically expressed in the PSM and when its expression is externally forced through optogenetics, the cells are entrained to this external rhythm(Shimojo et al., 2016). Notch signalling ligands, receptors and modifiers are also included in the set of oscillatory expressed genes(Dequeant et al., 2006; Matsuda et al., 2020). These experimental observations ultimately show that during the gene expression cycle, there is a phase interval in which the cell would be a signal sender, having Delta ligands, and another phase interval in which it would be a receiver, having Notch ligand properly modified to respond to Delta ligands from neighbouring cells. Hence, the communication is phase gated.

Here, we present a coupling mechanism based on unidirectional and phase-gated communication that reflects the Notch signalling directives and its dynamic employment by the PSM cells. It is based on the ability of oscillators to either send a signal, or respond to it, at specific and non-overlapping phase intervals as they cycle (Fig. 1a). Accordingly, we refer to this communication mode between oscillators as “walkie/talkie” coupling. Upon interaction of a signalling oscillator with a receiving oscillator, only the oscillator at the receiving end has the ability to respond. The response can range between two extremes. On one end there is complete stop of cycling (brake response) and on the other end extremely fast movement forward (accelerator response) (Fig. 1b).

**Figure 1.**
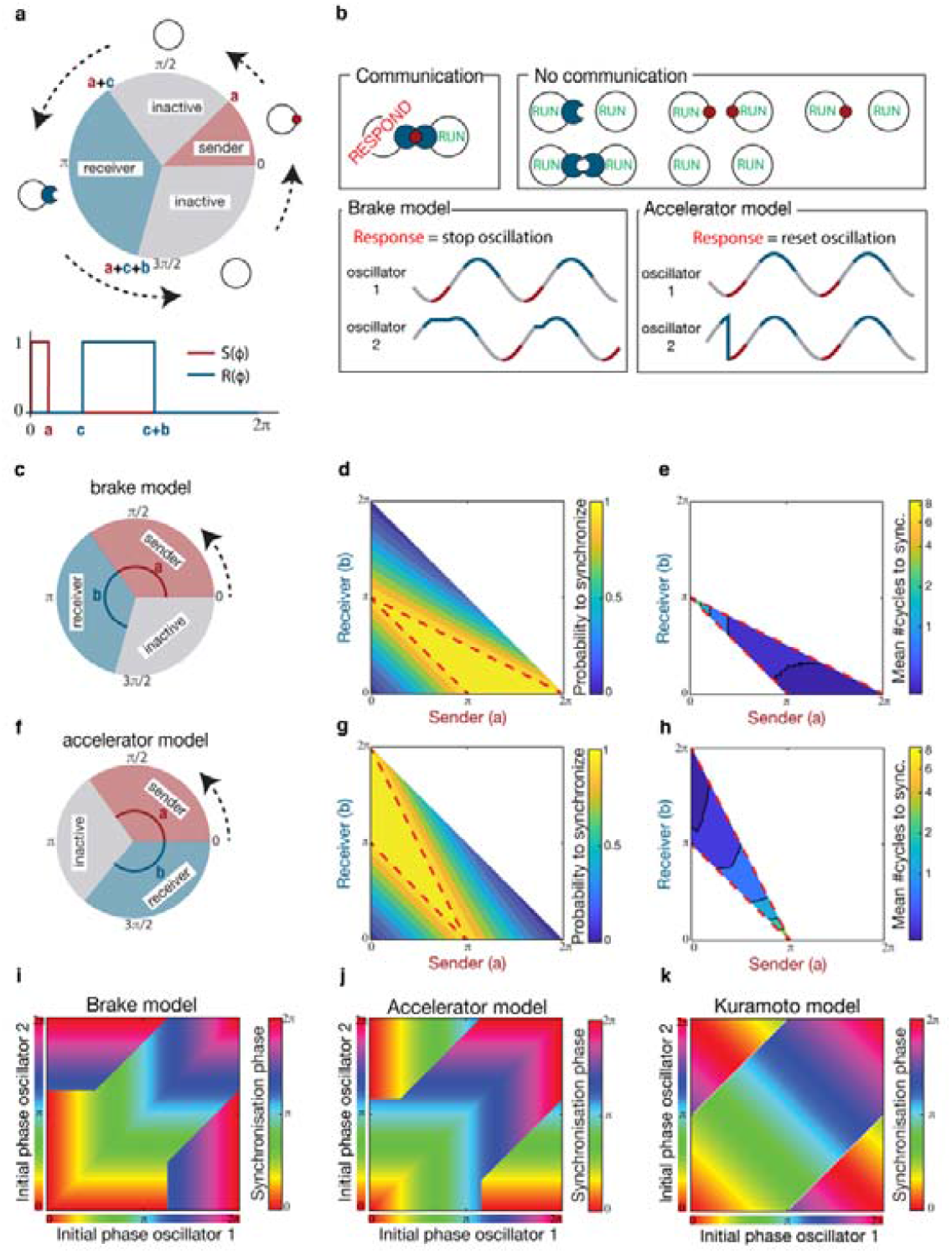
A coupling mechanism based on phase-gated and unidirectional communication synchronises two oscillators. **a**, During its periodic cycle, an oscillator goes through intervals when it either sends or receives a signal. Cycling is always counterclockwise. **b**, When an oscillator that sends a signal encounters one that is able to receive, efficient communication takes place and the receiver halts or accelerates its pace. All other encounters are not productive. **c**, Conditions for efficient synchronisation under the brake response require the signal receiving interval to immediately follow the sending one. **d**, Plotting the probability of synchronisation under brake response for all combinations of sending and receiving interval durations, a domain (highlighted by red lines) is identified where synchrony happens for all initial oscillator phases. **e**, In this domain we can plot the duration, in number of period cycles, until synchrony is reached, and this is below 2 cycles for the great majority of conditions. **f**, The receiving interval must precede the sending one when the responding oscillator accelerates. **g**, Plotting the synchronisation efficiency under the accelerator response for different durations of signal sending and receiving, a domain for certain synchronisation emerges that is a reflection along the diagonal of the brake response. **h**, Synchrony in this domain is also reached in less than 2 cycles for the majority of conditions. **i**,**j**,**k**, When plotting the synchronisation phase as a function of the initial phases of the interacting oscillators under the brake (**i**) or accelerator model (**j**), lines (isophases) parallel to the axes indicate that one of the initial phases entrains the other. For the Kuramoto model (**k**), such lines (isophases) are diagonal, indicating a compromise phase is reached between the two oscillators.

## Synchronizing two oscillators

In order to address whether and under which conditions the “walkie/talkie” coupling brings oscillators in-phase, we mathematically formulated a phase oscillator model (Supplementary Information). The only parameters are the duration of the cycle interval during which the oscillator sends a signal (a), the duration for the interval during which it can receive the signal (b), and the gap between them (c). The conditions for the synchronisation of two oscillators were found analytically and reveal mirror symmetries between the two opposite, extreme responses. For the brake response, the receiving interval has to follow the signalling interval (Fig. 1c), with a bias towards situations where signalling interval is larger than the receiving one (a>b) (Fig. 1d). For the accelerator response the order between signalling and receiving intervals has to be inverse, with the receiving interval preceding the signalling one (Fig. 1f). Additionally, the combinations of signalling and receiving interval durations are biased towards larger receiving intervals (a<b) (Fig. 1g). These findings can be intuitively understood, since in the brake model the responding cell has to be ahead, stop and wait in a small interval for a signalling cell to reach it. In contrast, for the accelerator model, the responding cell has to be lagging in a big interval from which it will be kicked forward to the signalling cell’s phase. Under these conditions, we see that synchronisation is achieved very fast, in less than two periods for the vast majority of conditions that ensure in-phase synchrony for the oscillators, for both brake (Fig. 1e) and accelerator response (Fig. 1h).

The synchronisation of two oscillators coupled with the walkie/talkie mechanism is characterized by the leading role of one of the two oscillators. The leader never adjusts its phase but forces the other coupled oscillator to follow and adjust in order to synchronise. The roles of the oscillators, as leaders and followers, depends on their initial phases (Supplementary Information). We have examined the behaviour of all possible pairs of oscillators under the “walkie/talkie” communication mechanism. If we extrapolate the oscillatory behaviour of the common pace to the time point that coupling was initiated, we can extract the phase on which the oscillators would settle if synchronization was achieved instantaneously (Suppl. Fig. 1). This defines the synchronization phase which can be understood as the phase on which the two oscillators land in order to start cycling in synchrony. Hence, an oscillator is a leader when its initial phase is the synchronisation phase. Under the “walkie/talkie” coupling, we see that there is always a leader (Fig. 1i, j). Importantly, the leader is a function of both phases, and it is impossible to say whether an oscillator will lead if the other coupled oscillator’s phase is unknown. The existence of a leader oscillator contrasts fundamentally with synchronisation under bilateral communication such as the Kuramoto model, for which a compromise phase is struck between the two oscillators’ initial phases (Fig. 1k).

### Synchronizing a group of oscillators

Subsequently, we extended our model to a field of oscillators connected with their immediate neighbours. We wanted to test whether “walkie/talkie” coupling is able to synchronise a group of oscillators arranged like the cells of the PSM. These cells oscillate with a period that varies along the anterior/posterior (A/P) axis(Tsiairis and Aulehla, 2016). In our model the intrinsic periods of the oscillators are randomly picked from a uniform distribution within the interval of measured periods in the PSM cells(Tsiairis and Aulehla, 2016) (Fig. 2a). We assume that a receiver oscillator responds if at least one of its direct neighbours is a sender oscillator (Fig. 2b). As cellular motility is observed in PSM cells(Bénazéraf et al., 2010) and has been important in understanding synchronisation dynamic(Uriu and Morelli, 2014), we allow neighbouring oscillators to stochastically swap positions with rate λ (Fig. 2b). We chose the rate λ such that a cell exchanges its position with one of its neighbours within 5-20 min on average, which reflects the measured movement of PSM cells(Uriu et al., 2010).

**Figure 2.**
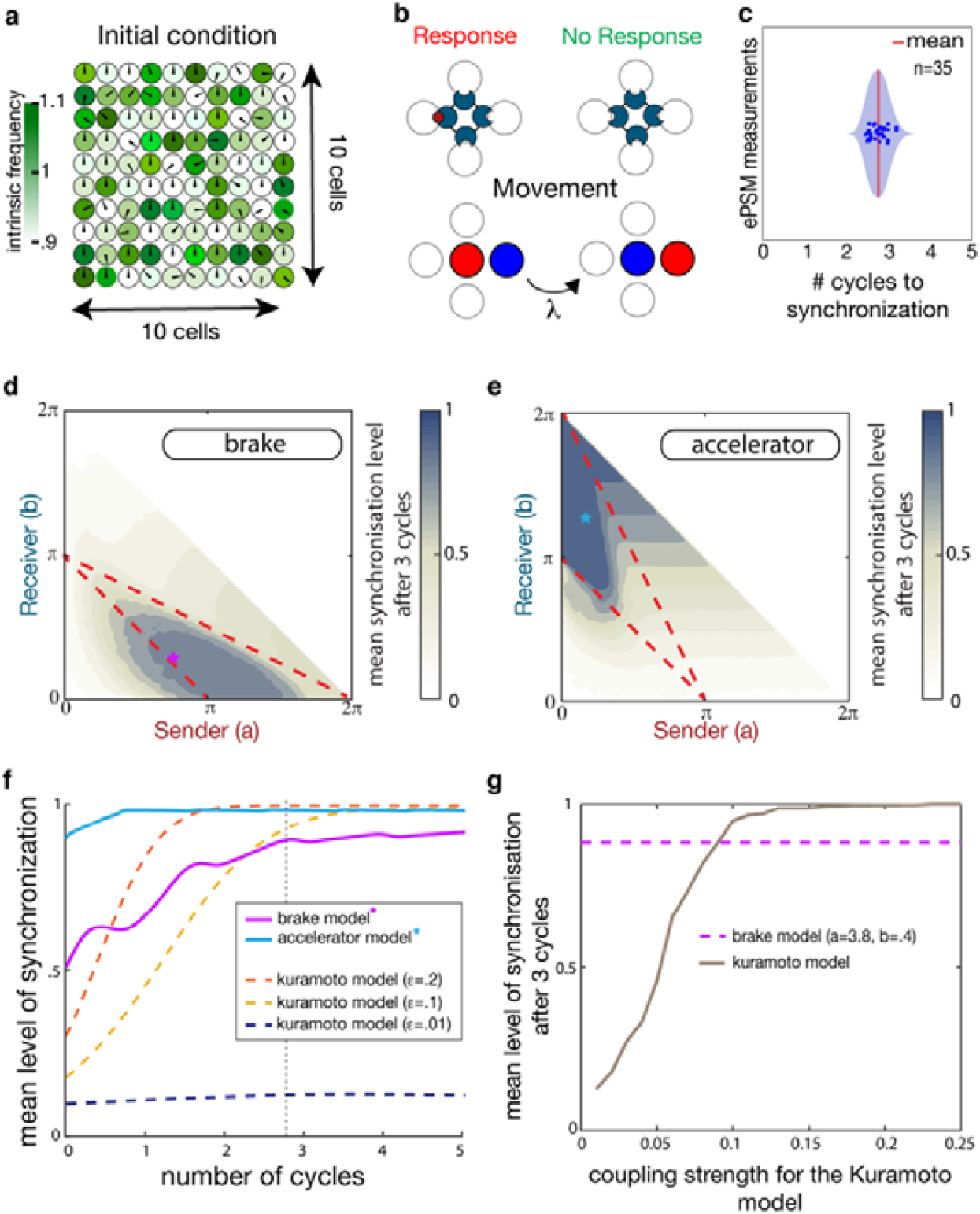
A group of locally coupled motile oscillators communicating via the “walkie/talkie” mechanism can be synchronised. **a**, Simulations of a field of 100 oscillators coupled with their immediate neighbours. Each oscillator has its own free-running period. **b**, For a receiving oscillator only one of the four neighbours being on the sending interval is sufficient for response, while motility is simulated with oscillators swapping places with a random neighbour at rate λ. **c**, Synchronised gene expression in the PSM is observed in less than 3 periodic cycles (2.76±0.19, n=35) after cells are randomly mixed. **d**, The average order parameter (z) that a group of oscillators reaches after 3 cycles under the walkie/talkie coupling, assuming the brake response, is plotted for all combinations of receiving and sending signalling interval. A domain where z>0.85 emerges mostly within the boundaries marking certain synchronisation for two oscillators (red line delimited). **e**, Similarly, assuming the accelerator response, the values of order parameter (z) are plotted against the combinations of sending and receiving intervals. A set of conditions emerges values of z approaching 1. **f**, The order parameter is plotted over time for a group of oscillators coupled via Kuramoto mechanism with different strengths (dashed lines) and is compared to the temporal evolution of order parameter for two selected conditions of the brake (purple star in d) and accelerator (blue star in e) “walkie/talkie” model. **g**, The order parameter reached after 3 oscillation cycles under Kuramoto coupling is plotted for different coupling strengths. The level of synchrony reached by “walkie/talkie” model (purple dashed line) is exceeded only for high coupling strength.

The level of synchronisation at any given time is given by the order parameter(Baker and Schnell, 2009; Kuramoto, 2012), a metric of how close are the phases of the coupled oscillators (Supplementary Information). For a large group of oscillators, the order parameter varies from 0, when the oscillator phases are uniformly distributed around the unit circle, to 1, when all the oscillators are perfectly synchronised. Synchronisation in the coupled gene expression oscillators of the PSM is observed in a time interval smaller than 3 periodic cycles (2.76±0.19, n=35) from the time coupling is enabled (Fig. 2c, Suppl. Fig. 2)(Tsiairis and Aulehla, 2016). Therefore, in the simulations, we calculate the level of synchronisation after 3 cycles. While for both the brake and the accelerator models a set of conditions allows for high mean level of synchronisation (up to .89 for the brake model and up to 1 for the accelerator) (Fig. 2d, e), this set is significantly smaller than for the two oscillator models. In addition, we see that cell motility increases significantly the level of synchronisation under the brake model, while it does not affect the synchronisation under the accelerator model (Suppl. Fig. 3). Improved synchronisation specifically for the brake model is also achieved as the number of connections per oscillator increases (Suppl. Fig. 3). Thus, it is possible that increased motility facilitates synchronisation by enabling more encounters for each of the coupled oscillators. When examining the level of synchronisation through time, we observe that both the brake and the accelerator models attain a high level already after the first cycle, whereas the Kuramoto coupling mechanism under the weak coupling conditions needs several cycles for a comparable level of synchronisation (Fig. 2f). Examining the synchronisation level at 3 cycles after coupling initiates, the brake model outperforms the Kuramoto mechanism under the weak coupling conditions and this is reversed only in the case of very strong coupling (Fig. 2g).

**Figure 3.**
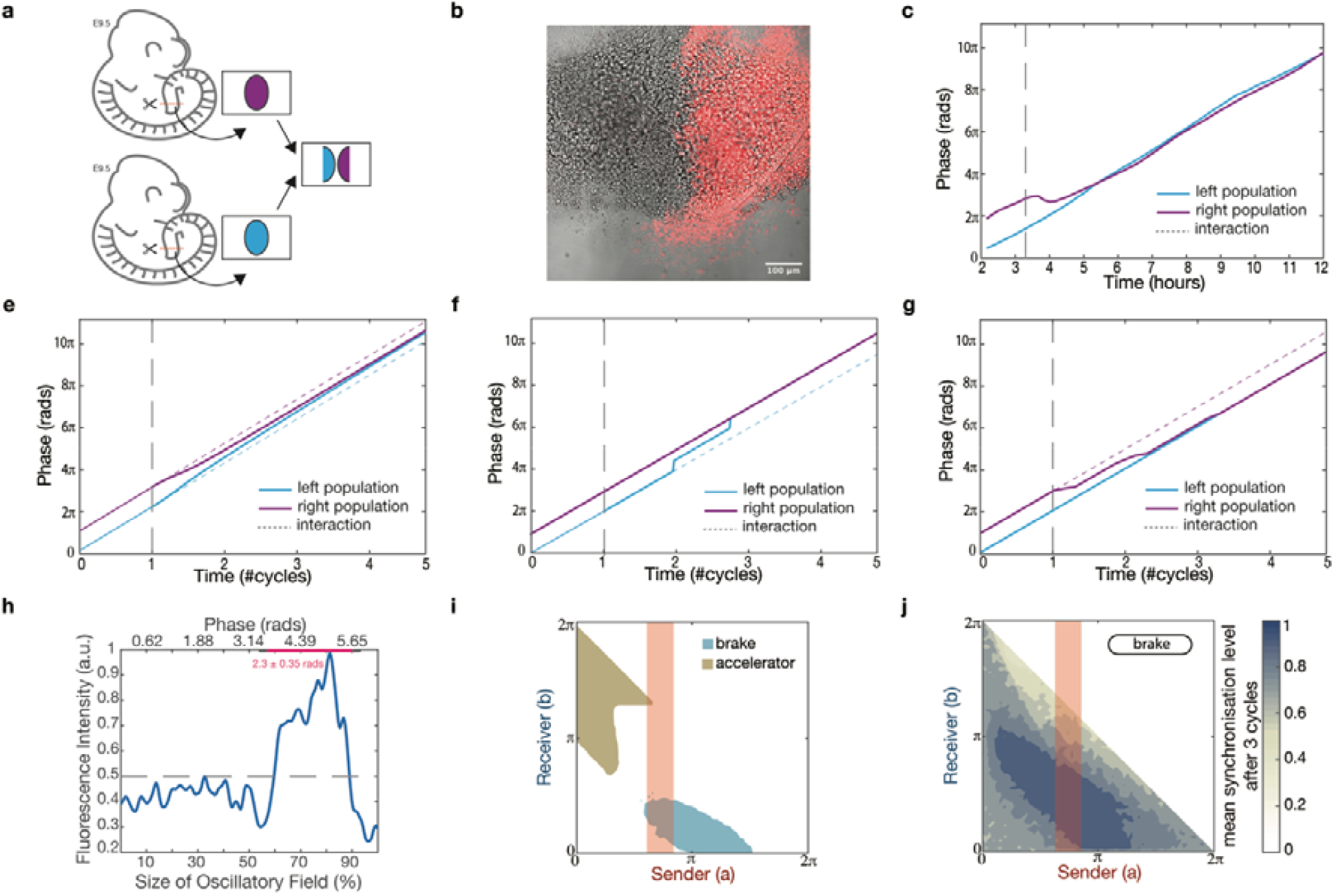
PSM cells halt their oscillation pace in order to synchronise. **a**, Tailbuds from different embryos are cultured in close proximity. **b**, As the tailbuds attach and grow on the fibronectin coated glass, the two populations of PSM cells collide and form a boundary. One population is labelled by the expression of H2BmCherry. **c**, The phase progression for the cells behind the boundary of two colliding PSM populations. One population slows down the rate of phase change (purple) while the other keeps cycling undisturbed (blue) (representative of n=6 independent experiments). The two populations encounter each other at the time point marked by the dashed line. **d**, Simulation of the phase progression for two populations coupled by Kuramoto mechanism shows that both populations’ phases drift from their original direction (broken lines) to converge over time. **e**, Simulating “walkie/talkie” coupling based on accelerator response leads only one of the populations to increase the slope of phase over time. **f**, When the response is of the brake type, the phase slope of the population that adjusts is decreasing. **g**, Quantification of Delta expression along the mouse PSM, where a full oscillation is unfolded in space, enables the calculation of signalling duration to be 2.3 ± 0.3 rad (n=10). **h**, The duration of signalling for the PSM cells is highlighted with red shade and overlaps with the set of conditions leading to synchronisation (z>0.85 after 3 cycles) under the brake response “walkie/talkie” (blue domain). **i**, The probability of two colliding populations to achieve synchrony is calculated through simulations and plotted for various durations of signal sending or receiving intervals. The red shade indicates the experimentally calculated range of signalling duration.

### Oscillation brake in the PSM coupling

The main difference between the brake and accelerator version of the “walkie/talkie” coupling mechanism lies in the way the phase of a receiving cell progresses, and we set out to record such phase progression in the PSM cellular system. Exploiting the mPSM assay(Lauschke et al., 2012), two tailbuds from different embryos, each containing cells that are synchronised, are cultured in proximity (Fig. 3a). As they spread on the fibronectin coated culture surface, the two populations collide with each other and form a clear boundary (Fig. 3b, Suppl. Fig. 4). Behind this boundary we can extract the oscillation phase and detect the changes as the two populations get synchronised and cycle in phase (Fig. 3c). While one population seems to behave as if there was no input, the complementary displays a sharp, temporary halt in the phase progression, that is never observed in free running mPSM cultures (Suppl. Fig. 4). To confront the experimental result to the model predictions, we reproduce the experiment in our model framework (Supplementary Information). A population that changes its rhythmicity according to the brake model displays a cycling break and the progressing phase temporarily slows down (Fig. 3e). In case the response is following the accelerator mode, the progressing phase displays sudden jumps forward (Fig. 3f). Applying the Kuramoto model, both populations will alter their pace in order to get aligned (Fig. 3g). It is clear that the experimentally observed behaviour correspond to the brake “walkie/talkie” coupling with one population cycling undisturbed, while the other slows down (Fig. 3c).

**Figure 4.**
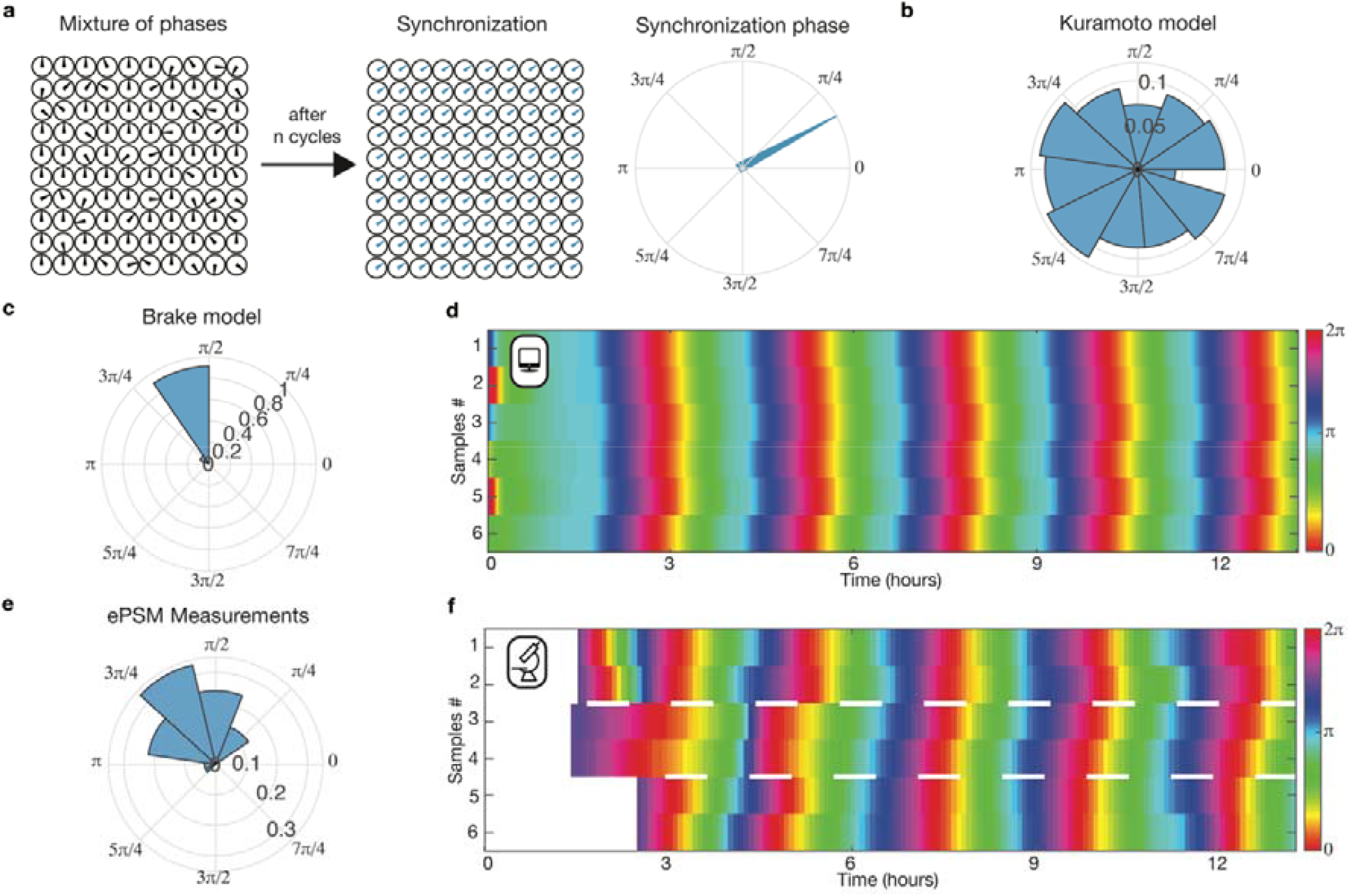
The coupling of PSM cells via “walkie/talkie” mechanism leads to a common phase after synchronisation. **a**, A group of oscillators with random initial phases get coupled at t_0_ and after n complete period cycles are synchronised with the same phase, called the synchronisation phase. **b**, Under Kuramoto coupling, simulations with different initial phases, indicate that the synchronisation phase is evenly distributed along all phases. **c**, Under the “walkie/talkie” coupling with brake response some phases are excluded from being the synchronisation phase. **d**, Different simulations of groups of oscillators appear synchronised when the phase of each group is plotted over time. **e**, Examining the synchronisation phase for different groups of PSM cells (n=54), they are distributed around a specific value (1.98±0.7 rads). **f**, The phase of different PSM cells that are in spatial and temporal (groups between white lines) isolation are plotted over time and apparent synchrony is observed. This is the result of the PSM cell groups reaching similar synchronisation phases.

As we have already theoretically identified conditions that enable a group of oscillators communicating via brake “walkie/talkie” coupling, we set out to examine a critical parameter reflecting the biological properties of the PSM. We focused on the easily measurable duration of signalling interval. This duration has to be sufficiently big for the brake response to be able to synchronise a population of motile oscillators (Fig. 2d). In the presomitic mesoderm cells, the signal sending state of cell is marked by high levels of Delta protein, the ligand for Notch signalling. We examined the relative duration of Delta protein presence, based on the fact that an entire oscillation cycle is unfolding at any given time along the length of the presomitic mesoderm(Lauschke et al., 2012) (Fig. 3h, Suppl. Fig. 5). Examining 10 samples, we calculated that the span of signalling is approximately 1/3 of the cycle (2.3 ±0.3 rads) (Fig. 3h), lying within the parameter region where the brake response efficiently synchronises a group of oscillators (Fig. 3i). It is noteworthy that if the signalling duration exceeds½ of the cycle would diminish the synchronization efficiency of the two colliding populations (Fig. 3j). Since the increased duration of signalling is generally beneficial for the brake response synchronization, we postulate an optimization of the Delta signalling to accommodate synchronization under multiple conditions.

### Synchronisation towards a specific phase

We next assessed whether the synchronisation of a group of oscillators coupled with the brake “walkie/talkie” mechanism is characterized by the leading role of one of the oscillators as shown above for two oscillators (Fig. 1i). We calculated the synchronisation phases for different random distributions of initial phases (Fig. 4a and Supplementary Information). When the cells are coupled through a Kuramoto mechanism, synchronisation phases cover the entire range of the cycle (Fig. 4b). This means that under this bidirectional symmetric coupling, any phase could be the phase on which a random initial distribution converges. Strikingly, under the brake “walkie/talkie” mechanism the prediction for the synchronisation phase value is very different, as it falls in a narrow interval around a specific phase that depends on the model parameters (Fig. 4c, Suppl. Fig. 6). As a consequence, the average trajectory of any group of well mixed oscillators will follow a predictable path to a specific synchronisation phase after less than 1 cycle (Suppl. Fig. 6). Subgroups of oscillators will converge to the same phase even if they are isolated from the others, adding robustness to the system (Fig. 4d). An actual reference oscillator doesn’t exist in the population. Nevertheless, the population behaves as if an apparent, phantom reference oscillator directs all the oscillators to a specific phase. The coupling mechanism guarantees this robust outcome.

We quantified the behaviour of the PSM cells expecting to observe this intriguing theoretical prediction. For this we exploited an assay where PSM cells are challenged to re-established synchronised oscillations. The tissue is mechanically dissociated in single cells that are aggregated into a clump, termed ePSM, where each cell encounters neighbours with variable phases. When cultured on fibronectin coated glass, the cells manage to reach a common pace and get synchronised in phase(Tsiairis and Aulehla, 2016). We measured the synchronisation phase for more than 40 such ePSMs (Suppl. Fig. 7). The resulting distribution is narrow and centered around a specific phase (Fig. 4e). Strikingly, as the model predicted, different groups of cells, that are cultured in spatial and temporal isolation, are synchronised in-phase despite this isolation (Fig. 4f). Our data are consistent with the brake walkie/talkie model and demonstrate that synchronisation of the PSM cells is robustly driven by unidirectional and phase-gated communication towards a narrow range of phases.

## Discussion

In order to understand the synchronisation mechanism behind the presomitic mesoderm cells we proposed a mechanism that captures explicitly the unidirectional and phase-gated communication through Notch signalling. This mechanism belongs to the broader category of pulse coupled oscillators(Winfree, 1967) and it contrasts the widely used Kuramoto mechanism on the communication pattern. It points to a different classification of coupling mechanisms according to this pattern with significant consequences. We demonstrate that such unidirectional and phase gating communication operates in the PSM and endows the cells with an overlooked property that provides robustness and predictability concerning the ultimate outcome of their synchronisation. Separated groups of well mixed cells synchronise towards the same phase. Such behaviour may have impact on the initiation of the synchronised oscillatory gene expression while the role of Delta as a trigger of an excitable oscillatory system(Hubaud et al., 2017) can provide a framework to incorporate the model deeper with the oscillatory mechanism.

The proposed model hinges on 2 main parameters, the duration of signal emission and reception, and thus provide a solid framework for future molecular investigations and analysis of PSM specific properties. Such a property is the gradient of period along the PSM cells which combined with the coupling mechanism can generate spatial patterns. Wave patterns appear in other such systems of pulse coupled oscillators(Ermentrout et al., 1984). Beyond the PSM, it is reasonable to expect the “walkie/talkie” communication coupling to underlie other synchronisation phenomena, where information is exchanged from a sender to a receiver, as is the case for chemical synaptic connections(Burns and Augustine, 1995). Finally, it could inspire approaches for fast and efficient synchronisation of engineered, artificial oscillators, because of its simple algorithm of few, clearly defined steps.

## Materials and Methods

### Mouse lines and timed matings

The LuVeLu transgenic reporter line(Aulehla et al., 2008) was used to visualize and quantify the oscillations in all experimental procedures. The R26-H2B-mCherry mouse line(Abe et al., 2011; Tsiairis and Aulehla, 2016) was exploited in combination with the LuVeLu in order to track the oscillatory activity of specific cell populations. Male animals of the mouse lines mentioned above were used in timed matings with wild-type CD1 mice to obtain embryos at E9.5 or E10.5 depending on the experimental needs. All experimental procedures were carried out in accordance with the Swiss animal protection laws and internal institutional guidelines.

### Emergent PSM (ePSM) cultures

Whole PSM tissues were collected from embryos originating from different pregnant females and pooled in groups of six or five. Single cell suspensions were achieved through mechanical dissociation as previously described(Tsiairis and Aulehla, 2016). Subsequently, the suspensions were passed through a 30μm filter (CellTrics). The cells were then centrifuged at 400rcf for 4min to form a pellet. Each pellet was then cut in four pieces. Every individual piece was placed alone in a well of a fibronectin-coated 8-well chamber slide (Lab-Tek). The pellets were cultured and imaged in culture medium (DMEM-F12 from Cell Culture Technologies containing 0.5 mM glucose, 2 mM glutamine, 1% BSA, 0.025/0.04 mg/mL penicillin/streptavidin) at 37°C and 5% CO2.

### Monolayer PSM (mPSM) cultures

E10.5 mouse embryos were dissected in dissection medium (culture medium containing 10 mM HEPES buffer). Tail-buds were surgically removed from the PSM of embryos as previously described(Lauschke et al., 2012). The tissues were then cultured on fibronectin-coated chamber slide (Lab-Tek) and imaged at 37°C and 5% CO2.

### Colliding mPSM cultures

E9.5 embryos were collected, and their tail-buds were surgically removed from the PSM ectodermal tissue was removed as much as possible using tungsten wire. Two pieces were then placed next to each other on a fibronectin-coated slide (μ-slide 8-well Ibidi chamber slide) and cultured in 37°C and 5% CO2. The positioning of the samples offered enough space for cell spreading and interaction between cells of the two samples. To allow fluorescence quantification from each sample individually, the R26-H2B-mCherry line was used. Comparable sample positioning, similar time of cell attachment, uniform spreading and viability were set as criteria to select samples for analysis.

### Live Imaging

Live imaging was carried out on a Zeiss LSM710 laser-scanning microscope. Depending on the experimental set up, the cultures were excited using an Argon Laser and a DPSS Laser at a wavelength of 514 nm and 561 nm respectively through a 20x plan apo objective (NA 0.8). Z-stacks of 3 planes at a distance of 5μm was scanned every 5 min (ePSM experiments) or 10 min (colliding mPSM experiments).

### Immunofluorescence

mPSM cultures were washed with ice-cold PBS and incubated on ice with fixative as described previously(Lauschke et al., 2012).The incubations with primary and secondary antibodies were performed at 4 °C overnight. Dll1 was detected with a 1:50 dilution of rat anti-Dll1 antibody (PGPM-1F9, MABN2284 from Sigma-Aldrich). After incubation the cultures were washed three times with 1% Triton-X-100, 10% FBS in PBS and then incubated with the secondary antibody (1:500 of Alexa Fluor® 488 goat anti-rat IgG) at 4 °C, overnight. Lastly, the samples were washed three times with 1% Triton-X-100, 10% FBS in PBS. Imaging was carried out on a Zeiss LSM710 system.

### Image processing

All data were acquired in 12bit, 512 × 512 pixels, 1.38μm/pixel. Preliminary image processing was carried out in Fiji. Maximum intensity projection was taken for every time-lapse. Hyperstacks were further processed by application of a Median filter (2.0 pixels radius) and a Gaussian Blur filter (σ = 10.0 μm). When needed, the nuclear signal coming from the R26-H2B-mCherry was used to generate a binary mask (based on a global threshold), which was subsequently used to collect the signal only from that particular sample.

### Time series analysis

The signals collected from Fiji were analyzed in MATLAB (Mathworks R2019b). Initially the signals were smoothed by applying a moving average of three time points. A detrended signal was extracted by taking the negative of the second derivative. In every derivation step, the signal was smoothened by moving average of three time points. Hilbert Transform was then used on the detrended signal to acquire the analytical signal as previously described and its phase was extracted(Lauschke et al., 2012). Linear fitting on the unwrapped phase was used to calculate the synchronisation phase at the time of centrifugation. The code used for generating the violin plot can be found at https://www.mathworks.com/matlabcentral/fileexchange/45134-violin-plot. All period measurements were carried out through Wavelet Transformation(Mönke et al., 2020).

## Acknowledgements

We thank Zoe Gruenig and the members of FMI animal facility for maintaining mouse colonies, as well as the Facility for Advanced Imaging and Microscopy for their support to the imaging required for this study. We are also thankful to Helge Grosshans, Katharina Sonnen and Catalina Chaparro for useful comments on the manuscript. Our work is supported by the Novartis Research Foundation.

## Author contributions

G.R. formulated the model mathematically, found analytical solutions, performed simulations, analyzed data and wrote the manuscript. G.M. performed experiments, analyzed data and wrote the manuscript. C.D.T. conceived and supervised the study, analyzed data and wrote the manuscript. The authors declare no competing interests.

---

**Suppl. Figure 1.**
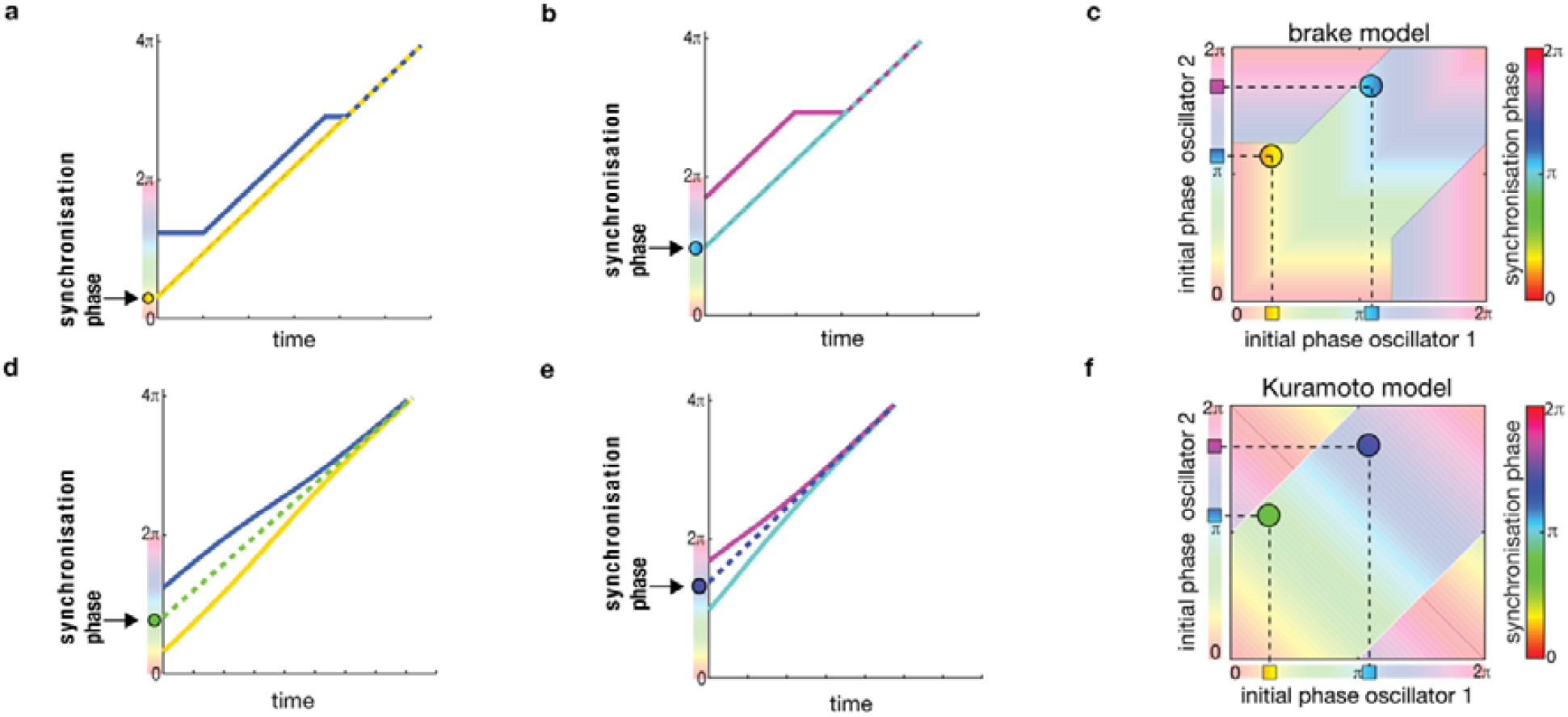
Generating the synchronisation phase plots for different coupling mechanisms. **a**, Two oscillators at different phases, one at blue and the other at yellow, get coupled at time 0 via the walkie talkie mechanism with brake response. Once the phases overlap and synchrony is reached the common phase is projected to time point 0 and it coincides with the yellow phase. **b**, A pair of coupled oscillators with different initial phases gets synchronised and the synchronisation phase coincides again with one of the initial phases, this time the light blue one. **c**, Combining different pairs we can generate a comprehensive plot with the synchronisation phase for all possible pairs. For the brake model the plot is generated after analytical calculation of the synchronisation phase for all possible initial conditions. **d**, When the oscillators from **a** are couple via Kuramoto mechanism, they get synchronised and the projection of the common phase to time point 0 falls between the initial phases. **e**, Similarly, the synchronisation phase for the pair of oscillators in **b**, under Kuramoto model is none of the initial phases and falls in between them. **f**, A plot for the synchronisation phase for all the combinations of initial phases is generated and is dominated by diagonal lines, in sharp contrast to the vertical and horizontal lines observed in **c**.

**Suppl. Figure 2.**
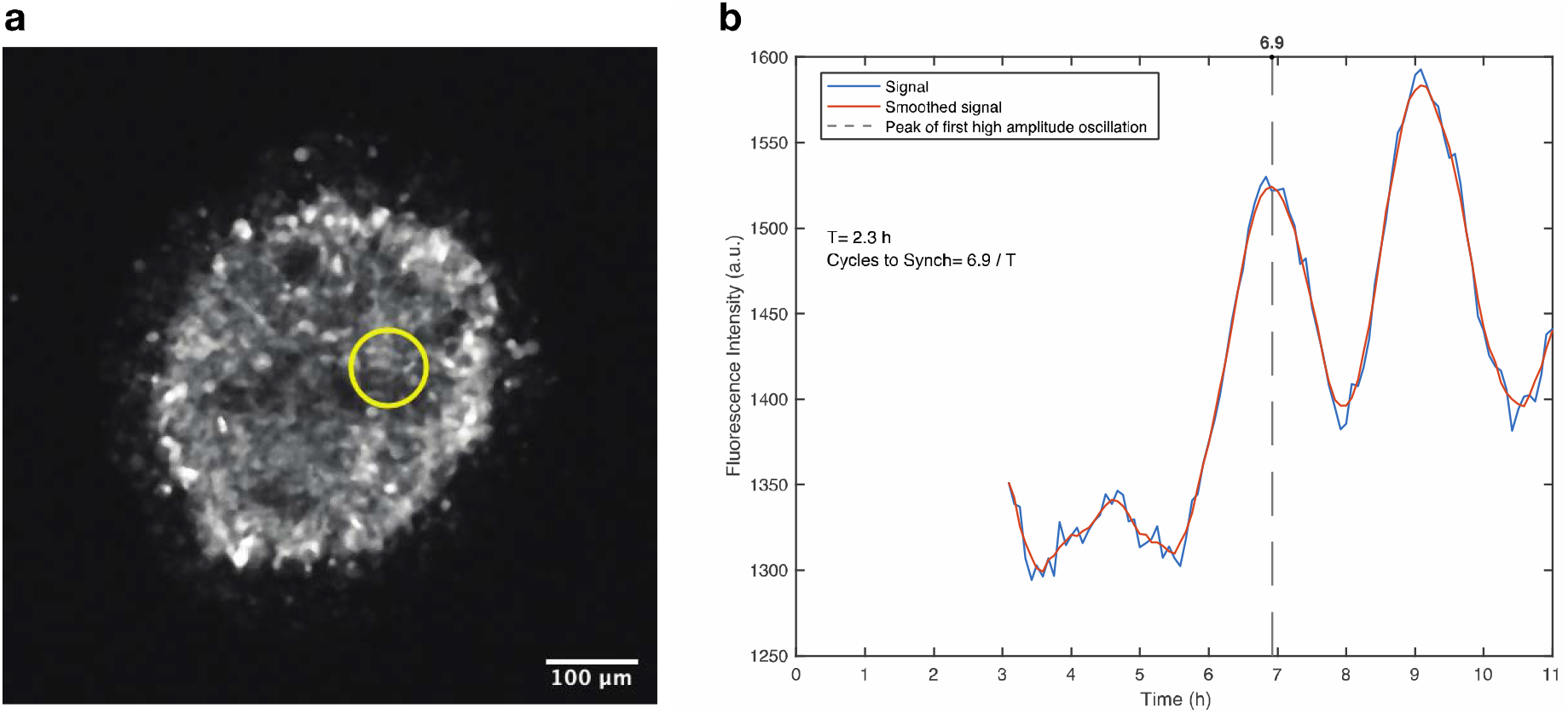
Estimating the time needed for synchronisation of PSM cells. **a**, Cells of the PSM are randomly mixed and left to synchronise in culture (ePSM) over a fibronectin coated glass. Visualizing the fluorescent reporter Luvelu through maximum intensity projection enables quantification of fluorescence intensity oscillations in the region marked by a yellow ring. **b**, The intensity of fluorescent reporter over time fluctuates and achieves the first high amplitude peak at the time marked by a dashed line. At this time coordinated gene expression of the cells has been achieved. At time zero the cells were centrifuged to form a clump, and this is the time at which communication via Notch signalling can be initiated. To express the time needed to achieve synchrony in cycles, the sum of minutes from centrifugation to the dashed line is divided by the period, T (for period measurements see Material and Methods section 1.7).

**Suppl. Figure 3.**
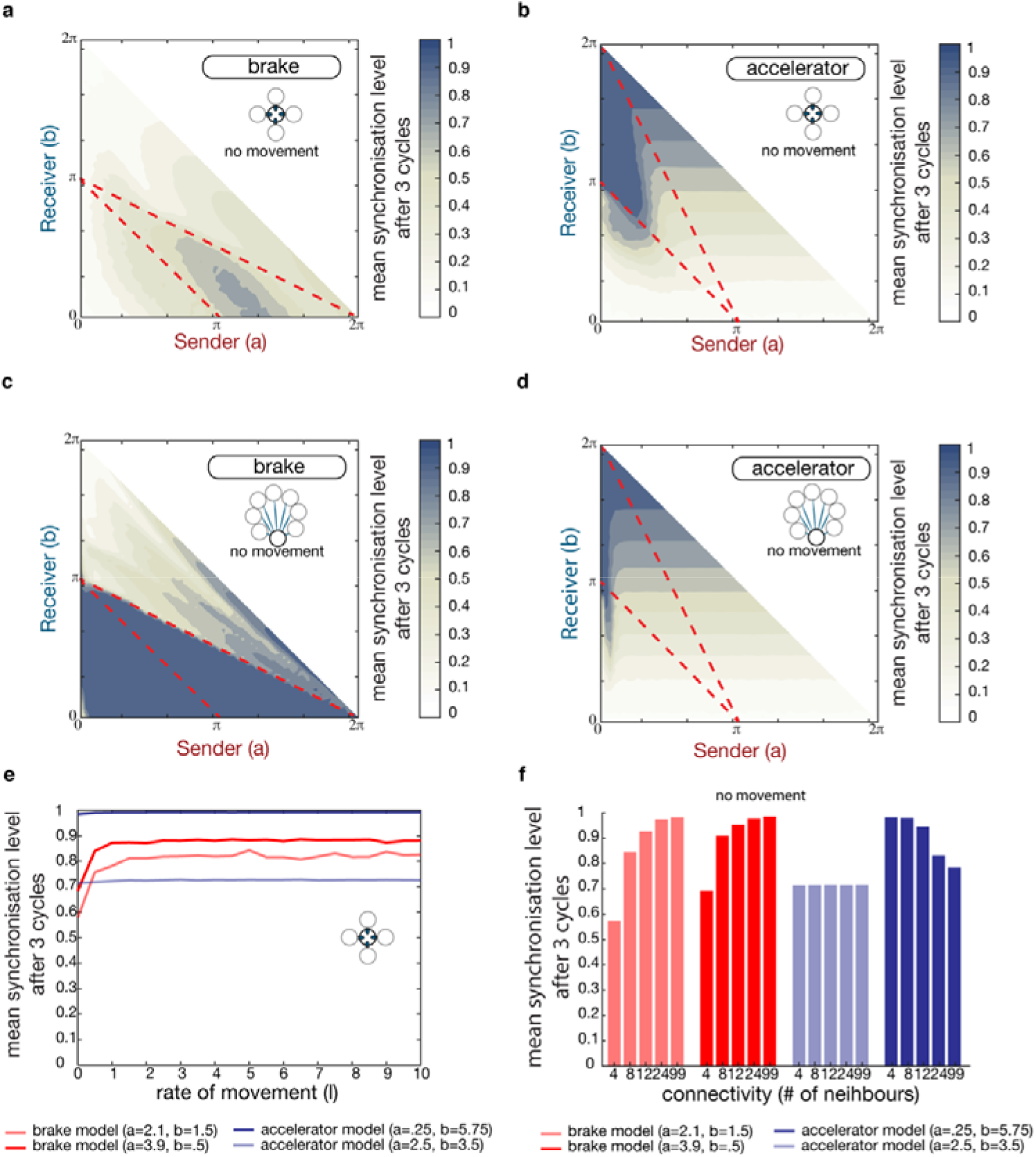
Examining the impact of brake and accelerator “walkie/talkie” coupling in the absence of motility. **a**, The mean order parameter after 3 cycles is plotted over the different combination of signal sending (a) and receiving intervals (b) for brake response. Each oscillator is interacting with only 4 neighbours and cannot move. **b**, Same plot for accelerator response. The connectivity is also limited to 4 neighbours. **c**, Plotting the mean order parameter after 3 cycles, when oscillators respond by breaking, and are connected to all others, shows an expanded set of conditions under which synchrony is reached. **d**, When the coupled oscillators respond by accelerating their pace, similar plots indicate that the parameter space under which synchronisation happens is shrinking. The accelerator model with all-to-all contacts and without motility. **e**, The mean order parameter after 3 cycles is plotted over different rate of movement (λ) for two set of parameters for the brake model (red) and two set of parameters for the accelerator model (blue). The number of neighbour contacts is set to 4. Motility has an impact on the brake model only. **f**, The mean order parameter after 3 cycles is plotted over different number of neighbour contacts for two set of parameters for the brake model (red) and two set of parameters for the accelerator model (blue); the motility is switched off (λ=0). The lack of motility improves synchronisation for the brake model but does not affect synchronisation of the accelerator model. Increasing the number of neighbour contacts enhances the synchronisation of the brake model but may imper the synchronisation of the accelerator model.

**Suppl. Figure 4.**
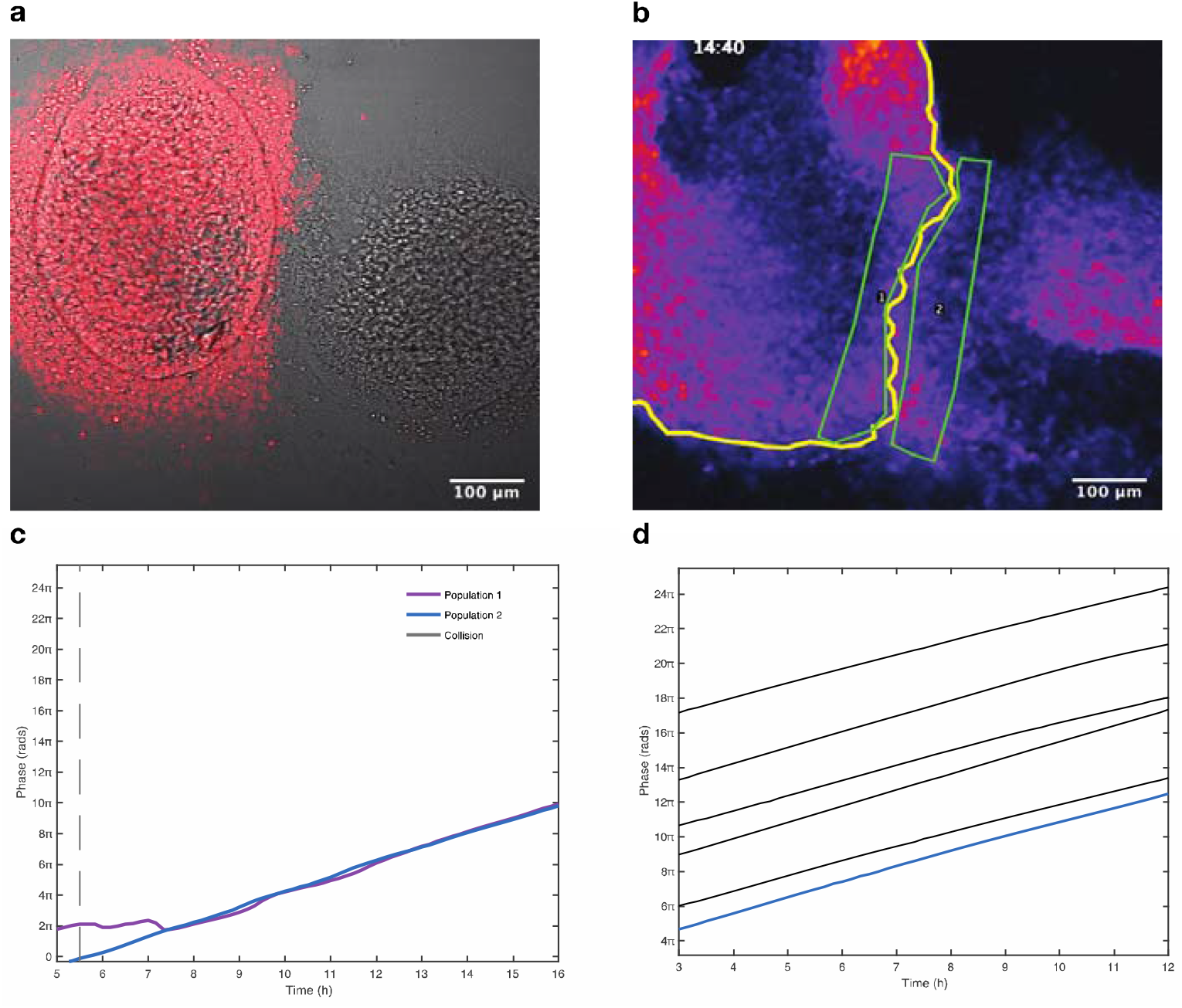
Quantification of the progressing phase behind the boundary of two colliding PSM cultures. **a**, Two tail-bud halves originating from different E9.5 mouse embryos are spreading and colliding with each other (MovieS1). The cell population on the left is expressing H2B::mCherry (Material Methods sections 1.1 and 1.4) enabling distinction of each culture. Cells from each tail-bud half stop spreading along a clear boundary and they do not mix. **b**, Regions of interest are highlighted with green borderline behind the line separating the two colliding cultures. The H2B::mCherry^+^ culture is marked by the yellow line. These are the areas for which the oscillating signal of the fluorescent reporter is collected and analysed. **c**, Plot of the progressing phase over time for the signal in the areas 1, and 2, as indicated in panel b. Population 1 exhibits a break response after the collision with population 2. Population 2 does not change its behaviour after the interaction with population 1. Dashed line indicates the point the two populations first meet. **d**, Plotting the phase progression of populations that haven’t collided with other cultures (black lines) allow to compare with the behaviour observed in c. In these populations there has never been observed a sharp kink in phase progression seen in the panel c for population 1 (purple line), while their behaviour is indistinguishable from the one of population 2 (added here with the blue line).

**Suppl. Figure 5.**
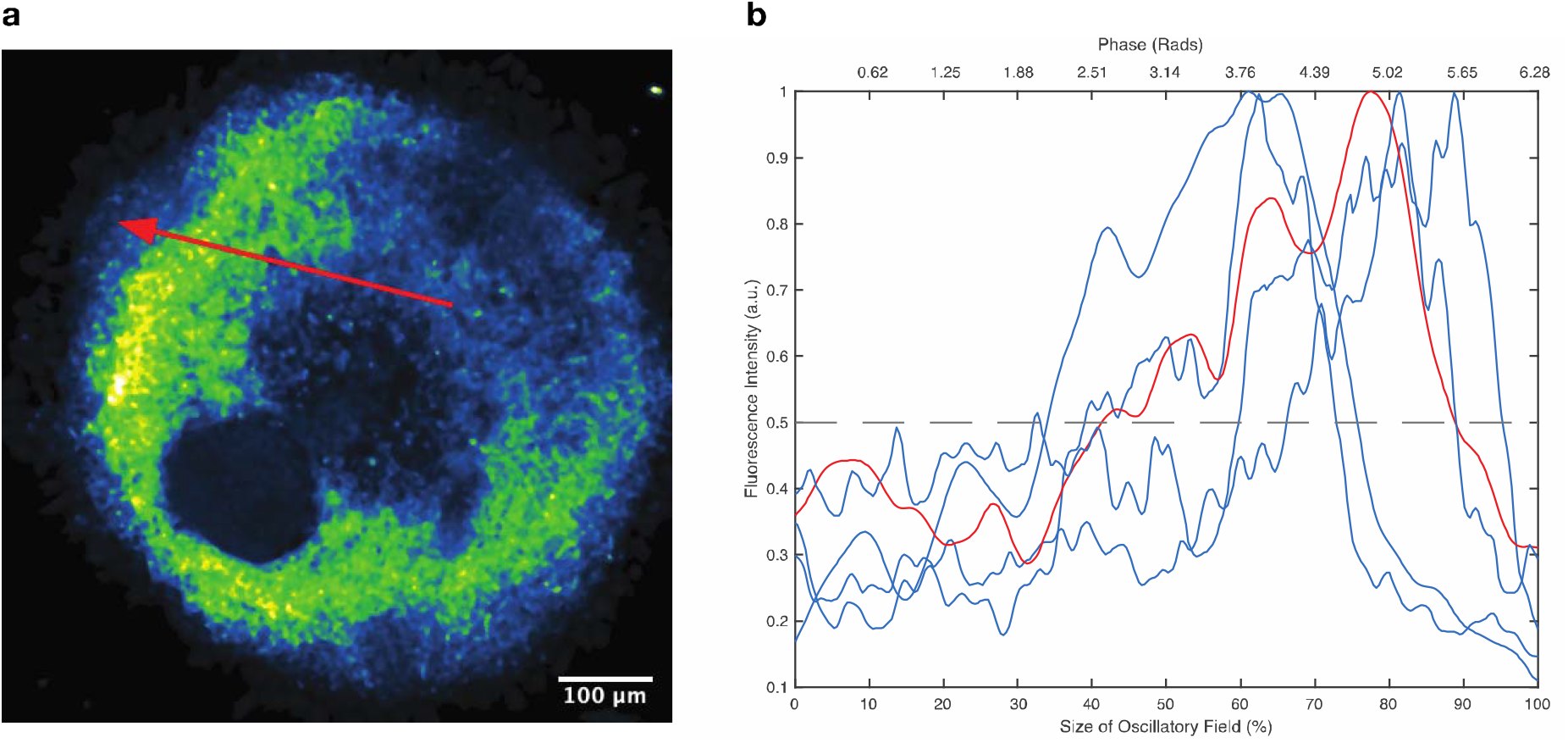
Quantification of the signalling duration, marked by presence of Dll1, during the oscillatory cycle. **a**, Representative immunofluorescence against Dll1 in a mPSM culture. The red arrow spans the culture from the centre to the periphery, along the path of transcription waves. **b**, The duration of Dll1 on the cycle is calculated by transforming the spatial domain into radians and using a threshold of 0.5. Measurements were performed in samples with similar Dll1 distributions (n=10). The profile along the arrow in panel a, is visualized with the red line. All profiles were normalized to the maximum intensity.

**Suppl. Figure 6.**
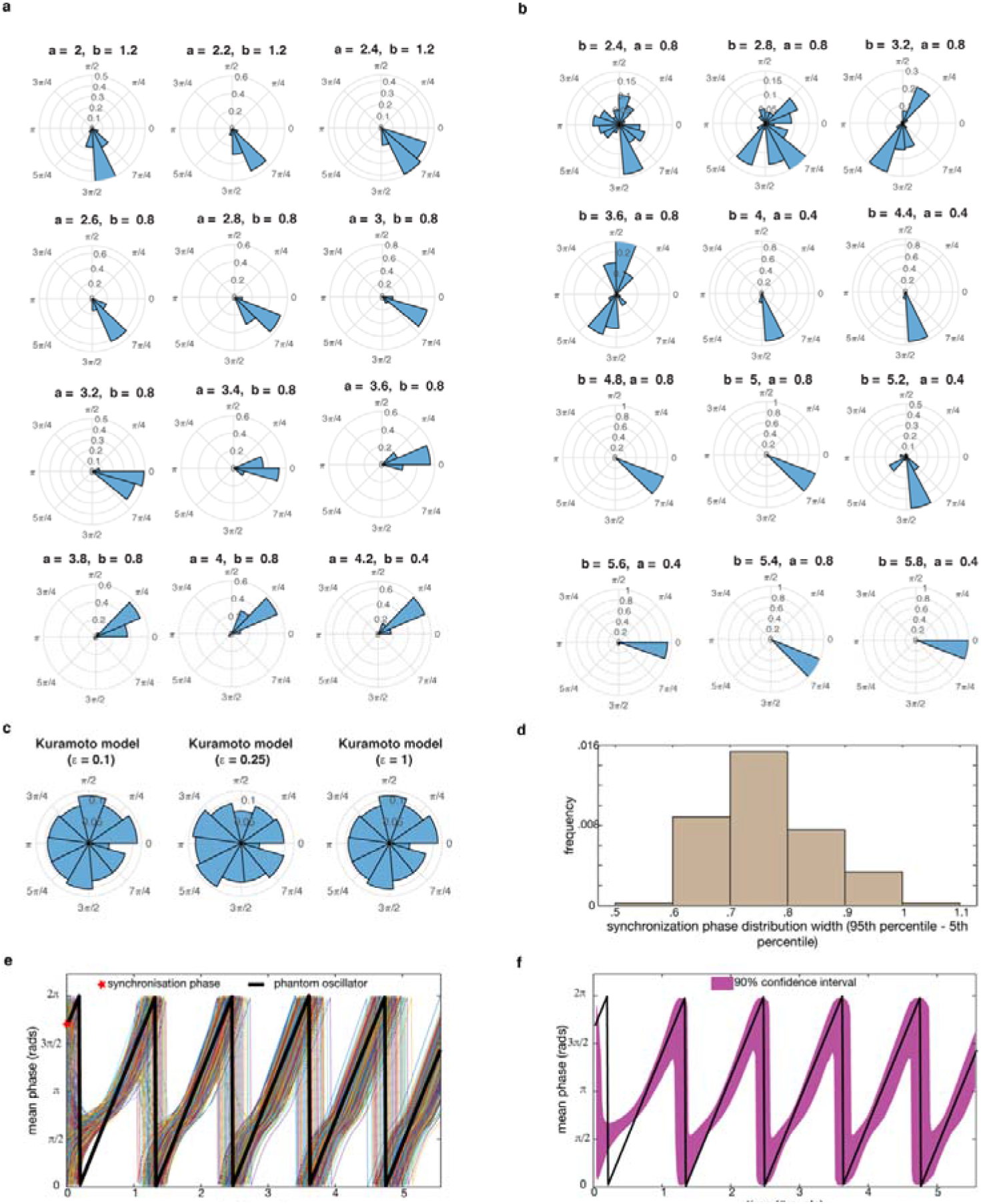
Distribution of synchronisation phases dependent on the coupling mechanism. **a**, Under “walkie/talkie” coupling with brake response, the distribution of phases under different durations of signal sending (a) and (b), always centres around a specific value. **b**, When the “walkie/talkie” response is of the accelerator type, we can identify conditions for (a) and (b) when we have synchronisation phase distribution with more than one peaks. **c**, For the Kuramoto coupling mechanism the synchronisation phase is uniformly distributed for all coupling strengths (ε) examined. **d**, To quantify the spread of the synchronisation phase distributions for the brake model, we calculated the interval length between the 5^th^ percentile and the 95th percentile of each distribution and plotted the histogram of these lengths. It is clear that the synchronisation phase under all examined conditions falls within a small section of the cycle. **e**, 1000 simulations of the brake model (a=2.1, b=1.5) were carried with differing initial phases, and the collected phase for all is plotted. Very quickly all converge to a similar behaviour, exemplified by a “phantom” oscillator that has the mean synchronisation phase as initial phase (black line). **f**, The fast convergence of all simulations towards the phantom oscillator (black line) is seen when we plot the interval of phases between the 5th percentile and the 95th percentile of the 1000 simulations.

**Suppl. Figure 7.**
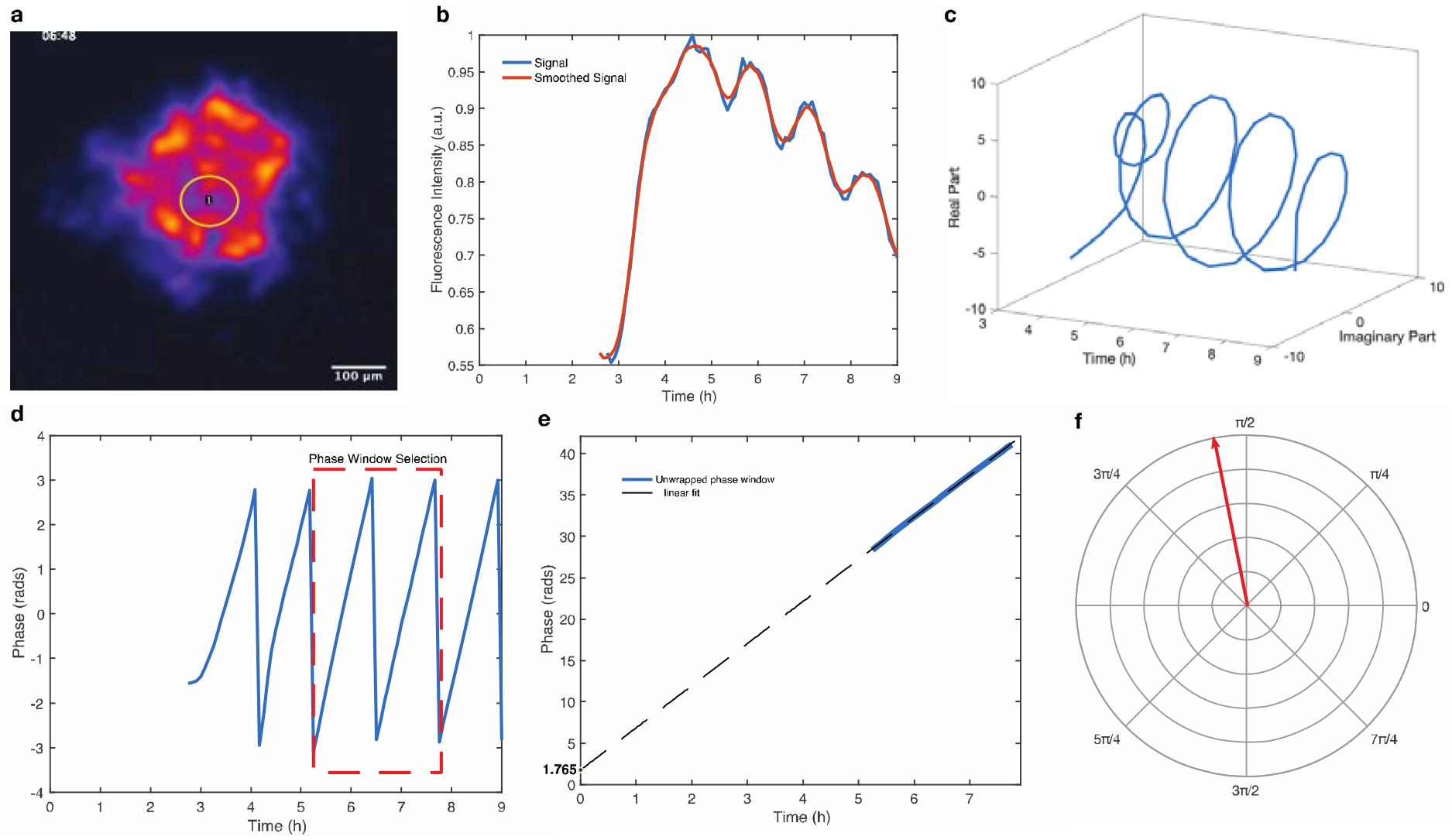
Calculating the synchronisation phase of presomitic mesoderm cell culture. **a**, Maximum intensity projection of an ePSM time-lapse experiment (see also MovieS2). **b**, Fluorescence Intensity profile for the region selected in panel a (yellow circular region of interest). Time zero corresponds to the time of centrifugation. **c**, Analytic representation of signal shown on panel b via Hilbert Transformation. **d**, Phase profile is acquired by calculating the angle of the analytic signal. A phase window of stable period oscillations is selected (in red). **e**, Plot of unwrapped phase over time for the selected phase window. Linear fit (dashed line) approximates how phase is changing over time. Using the linear fit, the synchronisation phase (phase at centrifugation) is projected at the time 0 as shown. The unwrapped phase (blue line) is expressed modulo 2π until the synchronisation phase is positive. The synchronisation phase in this particular case is 1.765 rads. **f**, Projection of the synchronisation phase for this sample on the cycle.

